# Metacognition and memory of emotional information: Judgments of learning predict the affectivity congruence effect in free recall

**DOI:** 10.1101/2020.01.07.897165

**Authors:** Marta Siedlecka, Agata Blaut, Borys Paulewicz, Joanna Kłosowska

## Abstract

Memory of emotional information often depends on the current mood and the dominant affective state. For example, studies show that people tend to recall emotional information of valence that is congruent with their affective traits. However, not much is known about whether this tendency is captured by metacognitive judgments of learning (JOLs). The aim of this study was to find out how people who score low or high on affectivity scales assess their memory of emotional material. We used a free-recall task with self-referential neutral, positive, and negative adjectives. The results show the affectivity congruence effect: the number of negative words recalled is related to affectivity; it increases with Negative Affectivity (NA) and decreases with Positive Affectivity (PA). Metacognitive assessment of future recall is also related to affectivity. Higher PA is related to higher JOLs for positive words and lower JOLs for negative words. Higher NA is related to higher JOLs for negative words and lower JOLs for positive words. The results suggest that metacognitive processes are sensitive to affective trait-specific memory bias.

---

Cognitive processing of emotional information often depends on the current mood or dominant affective state. For example, people suffering from anxiety display an attentional bias towards threatening sources of information (Bar-Haim, Lamy, Pergamin, Bakermans-Kranenburg, & Van Ijzendoorn, 2007; Cisler, & Koster, 2010; Mogg & Bradley, 1998; Williams, Mathews, & MacLeod, 1996) and depressed individuals recall more negative or unpleasant words and autobiographical memories than controls (Blaney, 1986; Gotlib & Joormann, 2010; Mathews & MacLeod, 2005; Matt, Vázquez, & Campbell, 1992). One of the best-studied phenomena of this kind is mood-congruent memory, which is the tendency to recall information of emotional value that is congruent with the current mood (Blaney, 1986; Bower, 1981; Eich & Macaulay, 2000; Eich, Macaulay, & Ryan, 1994). Mood-congruent memory bias has been observed in depressed, depression-prone participants, and in healthy participants in natural or experimentally induced moods (Blaney, 1986; Eich & Macaulay, 2000; Eich, Macaulay, & Ryan, 1994; Gotlib & Joormann, 2010; Mathews & MacLeod, 2005; Matt et al., 1992; Mayer, Cormick, & Strong, 1995; Parrott, 1991). Moreover, studies have also revealed the relation between emotion-related personality traits and memory bias: people scoring high in extraversion and positive affectivity questionnaires recall more positive memories, while people scoring high in neuroticism and negative affectivity tend to retrieve negative memories (MacLeod, Andersen, & Davies, 1994; Seidlitz & Diener, 1993; Rusting, 1999). Although biased memory of emotional stimuli has been observed in both clinical and non-clinical populations, not much is known about whether such bias is captured by metacognitive processes. The aim of this study was to find out how people characterized by different affectivity traits assess their memory performance in conditions in which the performance itself is biased.

Negative Affectivity (NA) and Positive Affectivity (PA) are mood-dispositional dimensions which reflect individual differences in emotionality. People scoring high on NA tend to be distressed and upset and they have a negative view of themselves, whereas those low on this dimension are relatively content, secure and satisfied with themselves (Watson & Clark, 1984). Individuals characterized by a high level of PA experience frequent and intense episodes of pleasant, pleasurable mood; in contrast, those who are low on the PA dimension report lower levels of happiness, vigour, and confidence. In theory, PA is independent of NA (see e.g. Watson, 2002). Theoretical concepts as well as the results of empirical work suggest that trait affectivity is related to the functioning of memory. The trait-congruency hypothesis states that certain personality traits predispose individuals to preferential processing of emotional information that is congruent with these traits (Gomez, Gomez & Cooper, 2002). According to Rusting (1998) this effect can be explained by Bower’s (1991) network theory of affect. Due to their susceptibility to experiencing different emotions, individuals characterized by different temperamental traits may develop different “emotion nodes”, which are cognitive networks composed of memories and cognitions related to particular emotions. Activation of a particular emotion node evokes emotion-related processes that, in effect, might bias cognitive processing.

Research suggests that extraverts, who are high on the PA dimension, have heightened emotional reactivity to positive mood induction procedures in comparison with introverts, who are low on PA. In contrast, people with high levels of NA (neurotics) seem to attend and respond more intensely to negative stimuli than stable individuals (Larsen & Ketelaar, 1989, 1991). People characterized by a high level of NA recall significantly more negative life events than those with a low level of this trait. This seems to be especially true for individuals who are not only high on NA but are simultaneously characterized by a low level of PA (Garcia & Siddiqui, 2009; MacDonald & Kormi-Nouri, 2013). Some authors (MacDonald & Kormi-Nouri, 2013) suggest that people high on NA tend to react more strongly to unpleasant events, and stress hormones have been shown to selectively enhance long-term memory of emotional stimuli (Cahill & Alkire, 2003; Cahill, Gorski, & Le, 2003).

Studies examining the relationship between affectivity and non-autobiographical types of memory also suggest the existence of trait-related memory bias. People high in extraversion have been shown to recall more positive and pleasant than negative and unpleasant material (Desrosiers & Robinson, 1992; Lishman, 1972), and reported happiness is related to more accurate recall of pleasant items (Matlin & Gowron, 1979). Research by Garcia and Siddiqui (2009) showed that in a task requiring participants to decide whether presented words were bold-typed in the story they had read earlier, all participants displayed a tendency to recognize more positive than negative words. However, people high on PA had a tendency to incorrectly identify negative words as not being presented previously. The authors interpret these results by stating that individuals with a high level of PA may be generally more prone to forgetting negative words than to forgetting positive words.

Is this memory bias captured by metacognitive assessments of memory performance? Metacognition is a term that refers to control and monitoring functions that enable people to assess their performance or knowledge and consequently choose the right strategy for dealing with a task. Metacognitive assessments have been extensively studied in memory research in which participants are asked to assess their own performance at different stages of a task. For example, in the study phase participants might be asked to predict their future recall of newly learned items (judgments of learning, JOLs, Arbuckle & Cuddy, 1969), and in the test phase they might be asked to assess the probability of recognizing the items they had not recalled (feeling of knowing, Hart, 1967) or to report their confidence that an item was recollected correctly (retrospective confidence judgments, e.g. Dougherty, Scheck, Nelson, & Narens, 2005). Metacognitive judgments are usually moderately accurate in the way that they correlate with the actual performance (Dunlosky & Nelson, 1994; Koriat, 1997; Mazzoni & Nelson, 1995), although they are susceptible to metacognitive illusions (Koriat & Bjork, 2006; Rhodes & Castel, 2008) or are blind to some cognitive effects, such as serial position effects in a memory task (Castel, 2008).

In this study we focus on judgments of learning that relate to the probability of future recall of each individual item (Mazzoni & Cornoldi, 1993; Nelson, 1993). Judgments of learning are thought to be based not only on multiple cues, such as general beliefs about one’s memory functioning and experience with similar types of tasks in the past (Hertzog, Dixon, & Hultsch, 1990; Mazzoni & Comoldi, 1993), but also on the properties of the items themselves (such as word frequency or concreteness, Koriat, 1997; Witherby & Tauber, 2017). JOLs are known to be impacted by the affective value of stimuli: they are typically higher for emotional than for neutral items (Hourihan, Fraundorf, & Benjamin, 2017; Nomi, Rhodes, & Cleary, 2013; Tauber & Dunlosky, 2012, Tauber, Dunlosky, Urry, & Opitz, 2017; Witherby & Tauber, 2018; Zimmerman & Kelley, 2010). There are two popular non-exclusive explanations for this effect: the first states that the distinctiveness of emotional information serves as a cue for predicting future recall; the second states that physiological arousal mimics the feeling of fluency or familiarity (Witherby & Tauber, 2018; Zimmerman & Kelley, 2010). The relation between JOLs for emotional items and performance has not been detected consistently: in some studies, emotional faces are associated with higher JOLs than neutral faces, but no differences in recognition are found (Nomi et al., 2013); in other studies, participants recall more emotional words, just as they predicted, but only in free recall and not in cued recall (Zimmerman & Kelley, 2010). Some studies suggest that arousal (not valence) biases judgments: higher JOLs are given to high-arousal items (that have neutral emotional value), even though no effect of arousal on actual recall or recognition was found (Hourihan et al., 2017).

Since there are discrepancies between memory performance and memory predictions when it comes to emotional stimuli, affectivity-related memory biases might not be reflected in judgments of learning. It is not clear whether people are aware of their memory biases; moreover, dominant affectivity might itself bias metacognitive judgments. Not many studies so far have focused on the relation between affective states and metacognition for non-emotional stimuli, although the few that did suggest that anxiety and neuroticism might be related to underconfidence (Colvin, Malgaroli, Chapman, MacKay-Brandt, & Cosentino, 2018), while positive mood might be related to overconfidence (Sidi, Ackerman, & Erez, 2018).

In this study we used emotional adjectives to test whether a person’s dominant affectivity is associated with memory bias and to find out whether this bias is reflected in metacognitive judgments. We used a free-recall task and free-recall judgments of learning to assess participants’ ability to predict the probability of future recall of each individual item (Mazzoni & Cornoldi, 1993; Nelson, 1993). We expected to observe affectivity-related bias in memory, namely that negative affectivity would be positively related to the number of negative words recalled and positive affectivity would be positively related to recall of positive words. Based on previous studies on JOLs and emotional stimuli, we could expect that JOLs would be higher for positive and negative words compared to neutral ones. On the other hand, dominant affectivity could differentiate between JOLS for negative and positive stimuli because memory-monitoring systems might have become sensitive to long-term memory bias. Therefore, we hypothesized that negative affectivity would be positively related to JOLs for negative stimuli, and positive affectivity would be positively related to JOLs for positive stimuli.

## METHODS

### Participants

One hundred and six undergraduate Jagiellonian University students from various faculties (89 female and 17 male; mean age = 23, SD = 8.2) completed the study in exchange for course credit. All participants gave their informed consent prior to the study. Because, to the best of our knowledge, there have not been previous studies on the relation between trait affectivity-congruent memory and JOL ratings and we did not have any preliminary pilot study results, we were not able to perform power analysis and estimate the required sample size. Therefore, we tested as many participants as our resources allowed^**1**^.

### Materials

#### *The Positive and Negative Affect Scales* (PANAS; Watson, Clark, & Tellegen, 1988)

PANAS assesses high-activation positive and negative states. The instructions of PANAS vary according to its state or trait version. We used the trait version with a general time frame: on five-point scales ranging from “very slightly” to “extremely”, participants rated how often in their daily lives they feel as was described by the statements. We used the Polish version of PANAS (Brzozowski, 2010).

#### Stimuli

The list of words used in the memory task consisted of 10 neutral, 10 negative and 10 positive adjectives. The word selection was based on previous research (Blaut, Paulewicz, Szastok, Prochwicz, & Koster, 2013) and the Nencki Affective Word List NAWL (Riegel, Wierzba, Wypych,Żurawski, Jednoróg, Grabowska, & Marchewka, 2015). The words were associated with mood and self-esteem (e.g., sad, useless, relaxed, happy). Because the Polish language has grammatical gender, we used masculine or feminine versions of the adjectives depending on the participant’s gender. Words of different emotional valence were matched for frequency, length and imageability.

### Procedure

Participants were tested in small groups in a computer laboratory. The experimental task was implemented in C++. We used LCD monitors (1280 × 800 resolution, 60 Hz refresh rate). After signing the consents, participants filled out the PANAS questionnaire. Then, the 30-trial memory task started. Each trial began with a 500-millisecond fixation point, which was followed by a 5-second presentation of a word and a JOL scale. Participants studied the word for 5 seconds and had to give JOLs before proceeding to the next item. The JOL scale was presented alongside with a question about how easy it would be to recall the presented word later. The scale was continuous but had 5 main sections: “Very Hard”, “Hard”, “Moderate”, “Easy”, and “Very Easy”. Participants were informed that they could choose different points within each scale category (with a mouse click). The study phase was followed by the distracting task, which lasted 3 minutes. Each trial of the distracting task began with a 1000 ms waiting period followed by a 500 ms presentation of fixation point. Then the words “right” or “left” were presented in the centre of the computer screen and participants were instructed to press the arrow key indicated by the word. Immediately after the distracting task, participants were asked to write down as many words from the memory task as possible within 3 minutes. At the end of the session, participants were questioned about what they thought was the goal of the study. No participants correctly guessed the exact hypotheses, although many noted that the items had different emotional values. At the end, participants were fully debriefed. The experimental session took approximately 20 minutes.

## RESULTS

The data were analysed using the R Statistical Environment (R Development Core Team, 2011). We used the lmerTest package (Kuznetsova, Brockhoff, & Christensen, 2017) to obtain approximate significance values for linear mixed models, and the lme4 (Bates & Sarkar, 2011) package was used to fit the logistic mixed models. Separate models with random valence effects associated with participants and random intercepts associated with items were fitted to estimate the effects of valence on JOL ratings (a linear mixed model) and recall probability (a logistic mixed model).

### Memory performance and judgments of learning: general results

Participants recalled more positive than negative words (*z* = 2, *p* = .04) and we did not detect differences either between negative and neutral words recall (*z* = 1, *p* = .3) or between positive and neutral words (*z* = 0.53, *p* = .6). JOLs were higher for positive words than for negative words (*t*(197) = 4.3, *p* < .001) and neutral words (*t*(197) = 7.6, *p* < .001). JOL ratings for negative words were higher than for neutral words (*t*(197) = 3.3, *p* < .001).

### Trait affectivity and memory performance

The results of PANAS revealed that mean PA score was 32 (SD = 15) and mean NA score was 20 (SD = 17). To test how the PANAS score is related to recall probability of different types of words, we fitted a logistic mixed model with fixed effects of valence, PA or NA nested within valence, and random intercepts associated with participants and items. We observed the trait affectivity-congruence effect for negative affectivity: NA was positively related to the recall probability of negative words (*z* = 2.2, *p* = .015, directional test). We did not detect statistically significant relation between PA and the recall probability of negative words (*z* = −1.9, *p* = .05), neither did we find significant effects of PA or NA scores on recall probability of neutral or positive words (|*z*| < 0.6, n.s.). However, NA was significantly more strongly related to the recall probability of negative words than to the recall probability of neutral words (*z* = 2.4, *p* = .014) and positive words (*z* = 2.6, *p* = .008), whereas PA was significantly more strongly related to the recall probability of negative words than to that of neutral words (*z* = 2.2, *p* = .03) but not positive words (*z* = 1.6, *p* = .1). The results are presented in Table 1 and Table 2.

**Table 1.**
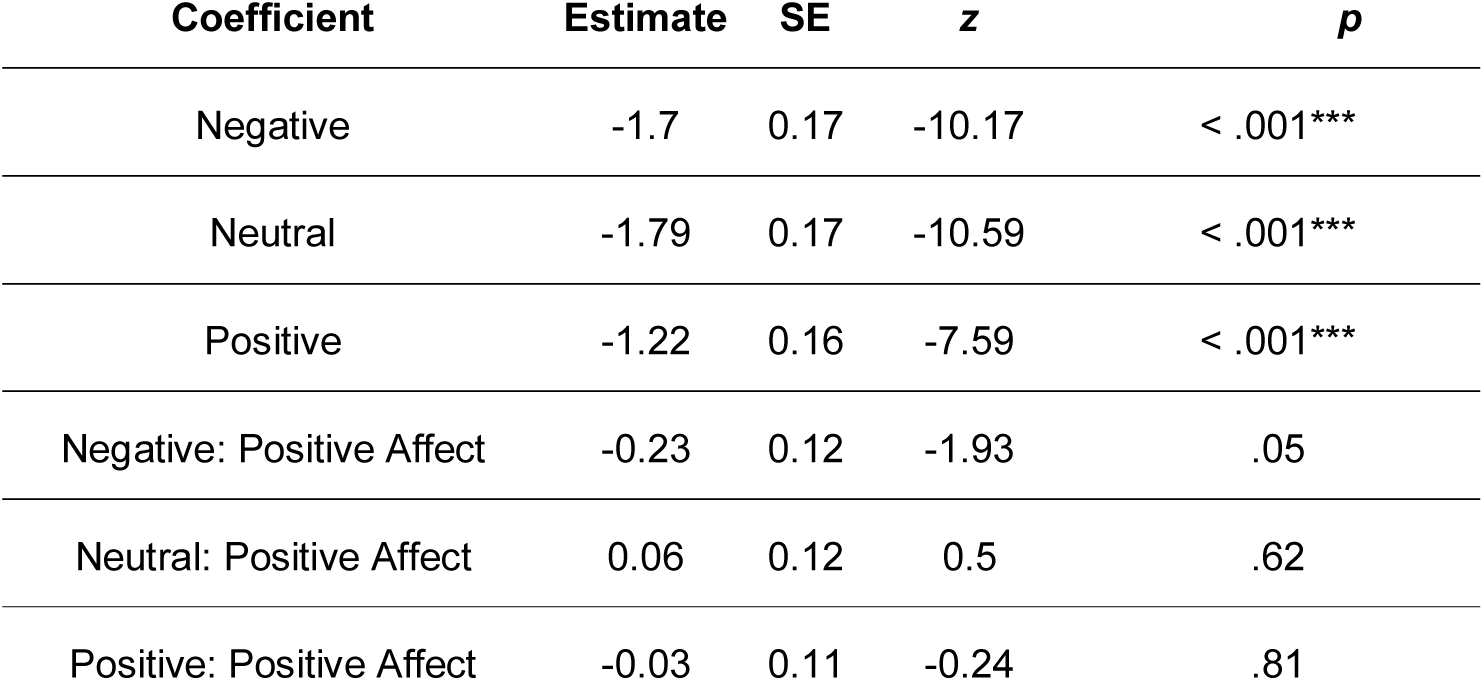
Model fit summary for mixed logistic regression with accuracy as dependent variable, fixed effects of both valence and Positive Affect, and random effects (intercepts) of subjects and words. Positive Affect scores were transformed to *z* scores to improve readability.

**Table 2.**
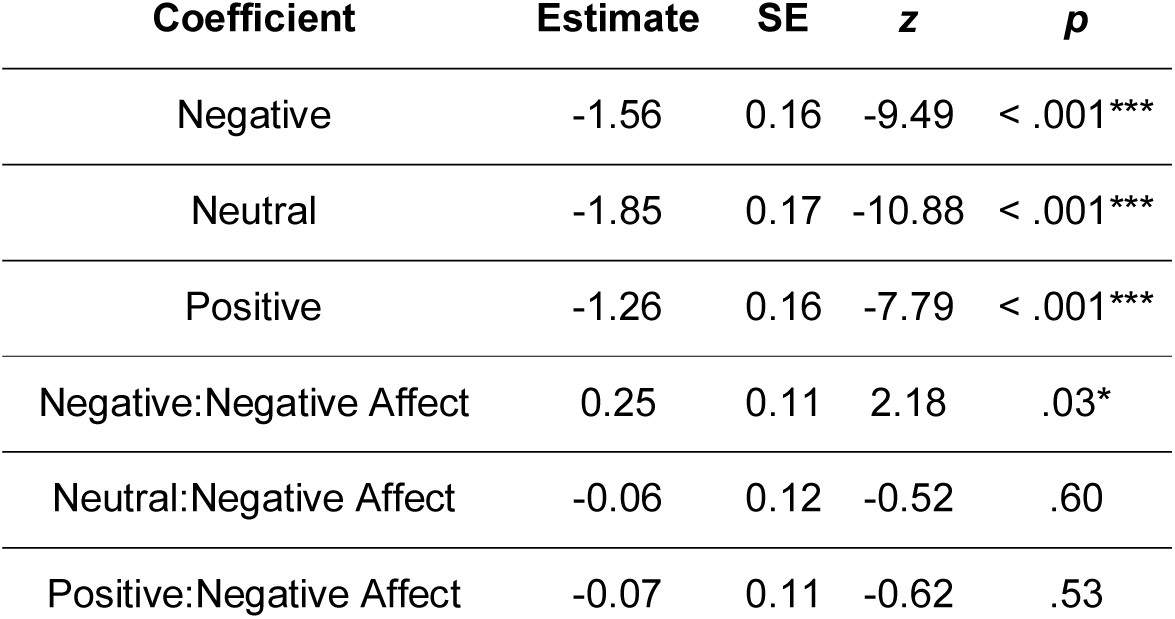
Model fit summary for mixed logistic regression with accuracy as dependent variable, fixed effects of both valence and Negative Affect, and random effects (intercepts) of subjects and words. Negative Affect scores were transformed to *z* scores to improve readability.

### Trait affectivity and judgments of learning

In order to test whether JOL ratings are related to PANAS score, we fitted a linear mixed model with fixed effects of valence, PA or NA nested within valence, and random intercepts associated with participants and items. PA was positively related to JOLs for positive words (*t*(254) = 0.84, *p* = .04) and negatively related to JOLs for negative words (*t*(256) = −0.03, *p* = .01). We found no relation between PA and JOLs for neutral words (*t*(254) = 0.84, *p* = .4).

NA was positively related to JOL ratings for negative words (*t*(219) = 2.5, *p* = .01) and was negatively related to JOL ratings for neutral words (*t*(217) = −3.0, *p* = .003). The effect of NA on JOL ratings for positive words was not significant (*t*(215) = −1.7, *p* = .09), although its direction (negative) reflected the analogous significant effect (positive) of PA. The results are presented in Table 3 and Table 4. A summary of all results is presented in Figure 1.

**Table 3.** Model fit summary for mixed linear regression with JOL rating as dependent variable, fixed effects of valence and Positive Affect, and random effects of subjects (intercept and valence) and words (intercept). Positive Affect scores were transformed to *z* scores to improve readability.

| **Coefficient** | **Estimate** | **SE** | **df** | ***t*** | ***p*** |
| --- | --- | --- | --- | --- | --- |
| Negative | 0.51 | 0.02 | 269 | 29.95 | < .001*** |
| Neutral | 0.46 | 0.02 | 268 | 27.09 | < .001*** |
| Positive | 0.62 | 0.02 | 268 | 36.54 | < .001*** |
| Negative: Positive Affect | -0.03 | 0.01 | 256 | -2.55 | .01* |
| Neutral: Positive Affect | 0.01 | 0.01 | 254 | 0.84 | .40 |
| Positive: Positive Affect | 0.02 | 0.01 | 251 | 2.03 | .04* |

**Table 4.** Model fit summary for mixed linear regression with JOL rating as dependent variable, fixed effects of valence and Negative Affect and random effects of subjects (intercept and valence) and words (intercept). Negative Affect scores were transformed to *z* scores to improve readability.

| **Coefficient** | **Estimate** | **SE** | **df** | ***t*** | ***p*** |
| --- | --- | --- | --- | --- | --- |
| Negative | 0.53 | 0.02 | 267 | 31.98 | < .001*** |
| Neutral | 0.44 | 0.02 | 266 | 26.65 | < .001*** |
| Positive | 0.60 | 0.02 | 266 | 36.18 | < .001*** |
| Negative: Negative Affect | 0.03 | 0.01 | 219 | 2.51 | .01* |
| Neutral: Negative Affect | -0.03 | 0.01 | 217 | -3.02 | .003** |
| Positive: Negative Affect | -0.02 | 0.01 | 215 | -1.7 | .09 |

**Figure 1.**
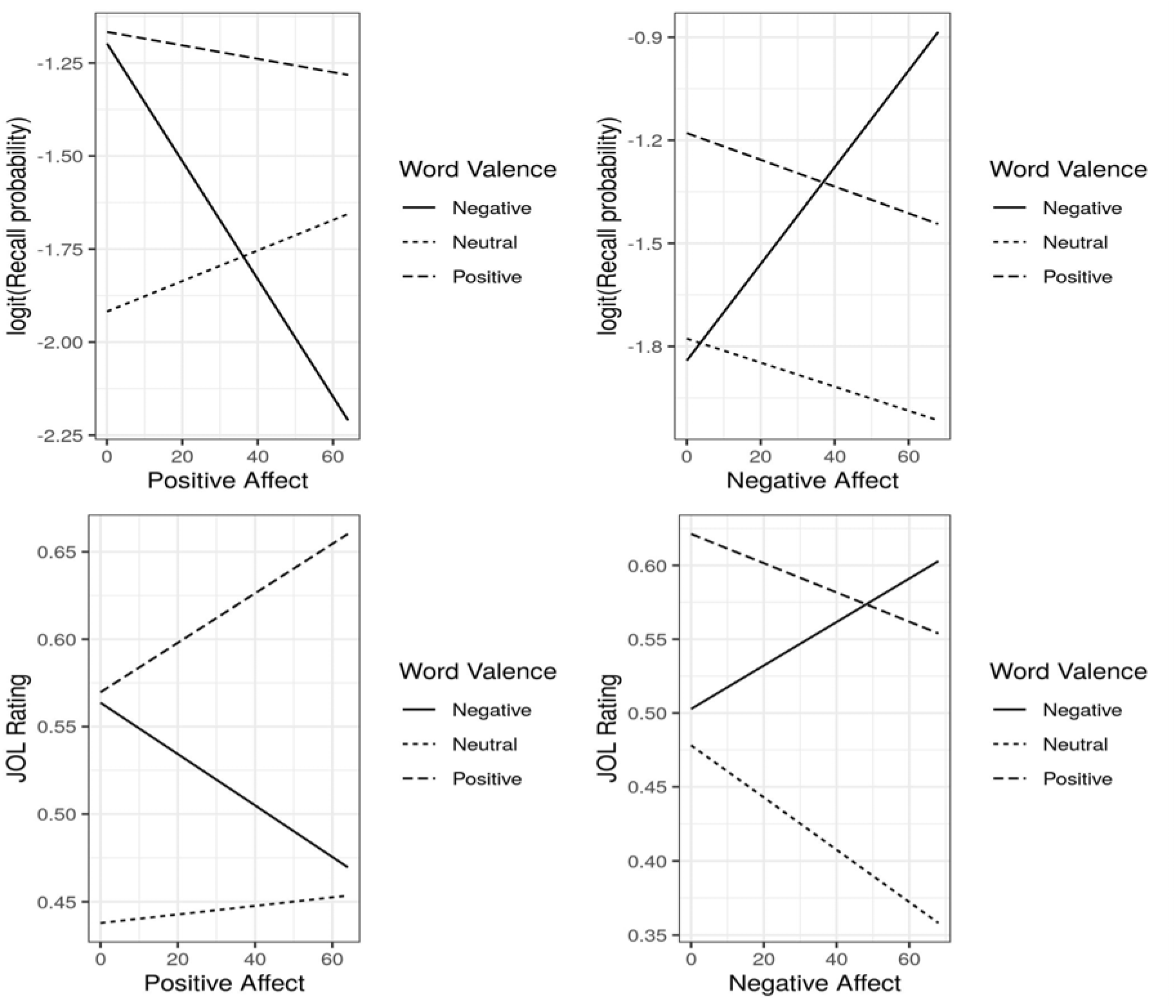
The relation between affectivity and Recall probability of differently valenced words (top panel) and between affectivity and JOL ratings of differently valenced words.

## DISCUSSION

In this study we tested whether the affectivity trait is associated with trait-congruent memory bias and whether this bias is taken into account when participants assess their memory. The results of the memory test show that participants generally recall more positive than negative and neutral words. However, the number of negative items recalled is associated with participants’ affective traits. The higher a person is on the NA dimension and the lower a on the PA dimension, the more negative words this person recalls, and these effects are significantly different from those observed for other word valences. Judgments of learning are generally highest for positive adjectives and lowest for neutral ones. However, affectivity traits are associated with qualitatively different ways of predicting future recall of words of different valence. People scoring higher on PA give lower JOL ratings for negative words and higher JOLs for positive words (no significant change for neutral items was observed). The NA dimension was related only to differentiating between negative and non-negative words: the higher the negative affectivity, the higher the JOLs for negative words and the lower the JOLs for neutral words.

The results support previous findings showing that trait affectivity is related to memory bias. However, participants typically learn easier and remember material consistent with their affective trait better than they remember material inconsistent with this trait (Garcia & Siddiqui, 2009; MacLeod et al., 1994; Rusting, 1999; Seidlitz & Diener, 1993). In this study we observed memory bias for items of negative valence: the number of negative words recalled correlated positively with the score on the NA dimension and negatively with the score on the PA dimension. Therefore, people higher on NA remembered more negative words than people lower on NA, and people higher on PA remembered fewer negative words than people lower on PA (see also: Garcia & Siddiqui, 2009; MacDonald & Kormi-Nouri, 2013). However, it is important to note that the trait-congruency effect is often calculated by comparing the number of positive and negative items retrieved, e.g. a person with high positive affectivity remembers more positive than negative items (e.g. Eich & Macaulay, 2000; Seidlitz & Diener, 1993). In our study we calculated the raw number of remembered items of each emotional value; this allowed us to assess the relation between this variable and the strength of trait affectivity.

The results of this study support the hypothesis that judgments of learning are generally sensitive to memory bias. Participants’ assessments of memory for negative words increased with NA and decreased with PA. However, this result could be interpreted in two different ways. Firstly, it seems that metacognitive processes, over time, adjust predictions of one’s behaviour in certain situations, or a person learns how his or her memory of stimuli that have a certain affective value works. If a person with high negative affectivity usually attends to and better remembers negative information, her metacognitive judgments will adjust to this cognitive bias. Studies show that people calibrate their metacognitive judgments in a relatively short time (during an experiment), for example, after receiving accuracy feedback (Miller & Geraci, 2011; Thompson, 1998) or after experiencing several study and test sessions (e.g. Castel, 2008; King, Zechmeister, & Shaughnessy, 1980). On the other hand, assuming that metacognitive functions not only monitor but also regulate information processing, it is possible that participants high on NA remember negative information better because this information is assessed by metacognitive processes as important. This metacognitive regulation might work differently for PA individuals as they might have a tendency to avoid or forget negative information (Garcia & Siddoqui, 2009). It is important in future studies to try to dissociate monitoring form the regulatory aspect of metacognition in order to better understand its role in memory bias and affective traits.

The current study might bring some insight into how JOLs for emotional stimuli are constructed. They are thought to be based on a number of cues, including subjective experience such as fluency or arousal when processing an item (Dunlosky et al., 2015; Koriat, 1997; Koriat & Ma’ayan, 2005). Contrary to other studies in which JOLs were higher for emotional versus neutral stimuli (Nomi et al., 2013; Tauber & Dunlosky, 2012; Zimmerman & Kelley, 2010), we observed that JOLs are sensitive to the valence of emotional words. The results of our experiment, which indicate that JOLs for negative words increase with increasing NA and decreasing PA, are in line with the hypothesis that high NA and low PA people are sensitive to negative self-referential information (Rusting, 1998). Therefore, these people might respond to such adjectives with high arousal, which in turn increases their JOLs (Witherby & Tauber, 2018; Zimmerman & Kelley, 2010). On the other hand, metacognitive judgments related to memorizing positive adjectives increase with positive affectivity (even though we did not detect a relation between memory of positive items and affective traits). Increased JOLs for positive words in high-PA participants could be explained as an effect of experienced fluency of processing trait congruence-related affective words. This interpretation would be even more interesting if we could argue that PA is indeed not related to the probability of remembering positive items and that the fluency itself inflates JOLs. Identifying conditions in which JOLs “track” memory performance and conditions in which the two variables dissociate would be very beneficial for deeper understanding of the mechanisms of metacognitive assessments and certainly needs further studying with a bigger sample size.

In conclusion, this is, to our knowledge, the first experiment to explore how metacognitive judgments reflect trait affectivity-related memory bias. The results show that metacognitive judgments predict trait-specific negative bias. Moreover, it seems that judgments of learning are sensitive to the trait-congruent emotional valence of stimuli. In future studies on this topic, it is important to study both the monitoring and regulatory functions of metacognition in order to understand how metacognitive processes track and control emotional cognitive biases.

The authors declare no competing interests.

## Footnotes

1 This study was a part of a larger project in which also depressive symptoms were measured (described in a separate paper that is in preparation). PANAS scores come from the same subjects but have not been described anywhere else.

